# Ultrastructural, metabolic and genetic determinants of the acquisition of macrolide resistance by *Streptococcus pneumoniae*

**DOI:** 10.1101/2023.12.27.573471

**Authors:** Xueqing Wu, Babek Alibayov, Xi Xiang, Santiago M. Lattar, Fuminori Sakai, Austin A. Medders, Brenda Antezana, Lance Keller, Ana G. J. Vidal, Yih-Ling Tzeng, D. Ashley Robinson, David Stephens, Yunsong Yu, Jorge E. Vidal

## Abstract

**Aim:** *Streptococcus pneumoniae* (Spn) acquires genes for macrolide resistance, MEGA or *ermB*, in the human host. These genes are carried either in the chromosome, or on integrative conjugative elements (ICEs). Here, we investigated molecular determinants of the acquisition of macrolide resistance.

**Methods and Results:** Whole genome analysis was conducted for 128 macrolide-resistant pneumococcal isolates to identify the presence of MEGA (44.5%, 57/128) or *ermB* (100%), and recombination events in Tn*916*-related elements or in the locus *comCDE* encoding competence genes. Confocal and electron microscopy studies demonstrated that, during the acquisition of macrolide resistance, pneumococcal strains formed clusters of varying size, with the largest aggregates having a median size of ∼1600 µm^2^. Remarkably, these pneumococcal aggregates comprise both encapsulated and nonencapsulated pneumococci, exhibited physical interaction, and spanned extracellular and intracellular compartments. We assessed the recombination frequency (rF) for the acquisition of macrolide resistance by a recipient D39 strain, from pneumococcal strains carrying MEGA (∼5.4 kb) in the chromone, or in large ICEs (>23 kb). Notably, the rF for the acquisition of MEGA, whether in the chromosome or carried on an ICE was similar. However, the rF adjusted to the acquisition of the full-length ICE (∼52 kb), compared to that of the capsule locus (∼23 kb) that is acquired by transformation, was three orders of magnitude higher. Finally, metabolomics studies revealed a link between the acquisition of ICE and the metabolic pathways involving nicotinic acid and sucrose.

**Conclusions:** Extracellular and intracellular pneumococcal clusters facilitate the acquisition of full-length ICE at a rF higher than that of typical transformation events, involving distinct metabolic changes that present potential targets for interventions.

## Introduction

*Streptococcus pneumoniae* (Spn), a Gram-positive opportunistic pathogen, is a leading cause of community-acquired pneumonia, meningitis, and otitis media^1, 2^. The treatment for pneumococcal disease has been challenged in recent years due to the rise of antibiotic resistance, including resistance to macrolides and β-Lactams in pneumococcal strains^3–5^. While the incidence of pneumococcal disease has decreased after the introduction of pneumococcal conjugate vaccines^6, 7^, vaccine escape strains are emerging as a public health threat due to serotype replace^7, 8^.

Despite causing disease, Spn asymptomatically colonizes the nasopharynx during childhood, forming biofilms (i.e., bacteria aggregates) and sharing this niche with several other streptococci species^9–13^. Pneumococci forming nasopharyngeal biofilms enter a dormant stage that slows down their metabolism thus allowing persistent colonization that often lasts for months^12, 14^. Persistent colonization is accompanied by a concomitant reduction of the expression of virulence factors, such as the capsular polysaccharide instead forming a biofilm structure made of extracellular DNA, proteins, lipids, and polysaccharides that facilitate asymptomatic carriage^10, 12, 13^. We and others have demonstrated that pneumococcal strains form biofilm consortia on abiotic surfaces as well as on human nasopharyngeal cells^15–18^. Furthermore, human nasopharyngeal cells, cultivated in a bioreactor system, facilitated the swift, nature-like acquisition of antibiotic resistance genes between two Spn strains with a high recombination frequency (rF)^15, 19, 20^.

Macrolide resistance in Spn is mainly attributed to modification of the macrolide target in the bacterial ribosome and to active efflux mediated by the <u>m</u>acrolide <u>e</u>fflux <u>g</u>enetic <u>a</u>ssembly (MEGA) element^5^. The most common mechanism altering the bacterial ribosome target involves the methylation of the 23S rRNA by methyltransferases encoded by *ermB* and rarely by *ermTR*^21^.

Point mutations in the 23S rRNA or L4 or L22 ribosomal protein genes *rplD* and *rplV*, respectively, have also been associated with non-susceptibility to macrolides^22, 23^. Another mechanism conferring macrolide resistance to Spn strains is the two-component (MEGA) efflux pump^24^. MEGA is encoded by a ∼5.5 kb, or 5.4 kb, element carrying the *mef(E)* and *mel* (listed also as *msrD*) operon^24, 25^. Spn strains carrying MEGA are resistant to 14- and 15-membered macrolides but they are susceptible to lincosamides and streptogramin B, known as the M phenotype^26^.

The *mef(E)*/*mel* genes and/or *ermB* are carried in large integrative conjugative elements (ICEs) of the Tn*916*-related family and Tn*5253*-related family, which also often carry the *tetM* gene for resistance to tetracycline^27–29^. The Tn*916*-related elements carrying MEGA inserted into *orf6* are termed Tn*2009*^28, 30^. Tn*916*-related elements were identified in macrolide-resistant Spn strains isolated in the late 1960s^31^ but their prevalence has significantly increased in the last few years. Invasive and nasopharyngeal Spn strains carrying Tn*916*-related ICEs, including Tn*2009* and other ICEs, have been identified in China, Italy, Venezuela, USA, Spain, and the UK^28, 29, 32, 33^. Moreover, ∼90% of emergent Spn vaccine escape strains of serotype 35B and 35D, isolated in Japan, carry macrolide and tetracycline resistance determinants in Tn*916*-related ICEs; of these 75% Spn strains carried MEGA in Tn*2009*^34^. Vaccine escape serotype 3 strains clonal complex (CC180) clade II, isolated in the USA and Hong Kong, carry *ermB* and *tetM* in a 36.7 kb Tn*916*-related ICE^35^.

In other Gram-positive species, the transfer of Tn*916*-related elements relies in the conjugation machinery (i.e., a type IV secretion system, T4SS), self-encoded in these ICEs^36^. The molecular mechanism by which ICEs are acquired by Spn strains has been debated for over 25 years. However, the most common hypothesis suggests that acquisition occurs via transformation^26, 28, 30^. This hypothesis was based on the observation that conjugative insertion of Tn*2009* or Tn*2010* in Spn strains were infrequent^28, 29^, that *in vitro* conjugation assays has been unsuccessful to generate transconjugants^26, 30^ or that the conjugative transference of Tn*916* or Tn*916*-related ICEs, when it has been observed, it has occurred at a low (<10^-6^) conjugation frequency^37^. In recent years, we developed in our laboratories a life-like bioreactor system that facilitates the natural transference of mutation-mediated resistance^15^, as well as macrolide resistance carried in ICEs among Spn strains^19, 20^. Using the bioreactor system, we demonstrated that the transformation machinery, specifically the Com system, facilitates the acquisition of MEGA, carried in Tn*916*-related elements, at a recombination frequency (rF) as high as 10^-3^. In the absence of either a functional Com system or a functional transformation apparatus, the rF of the acquisition of Tn*916*-related ICEs fell to <10^-7, 19^.

Competence development is a physiological state that is tightly regulated by certain environmental cues and cellular determinants. Although acquisition of antibiotic resistance naturally occurred whole strains colonize the human host, the host cell determinants are largely unknown. These host cells-derived molecules are mimicked in the bioreactor system, as pneumococci becomes naturally competent, release extracellular DNA and acquire DNA at a high recombination frequency when cultured in the bioreactor^15, 19^. This occurs even in the absence of antibiotic pressure, and within a few hours of incubation. At present, important molecular events, cellular factors, and the cellular dynamics of ICE transfer among Spn strains have remained unexplored.

In a recent study, we isolated and whole genome sequenced Spn strains from pneumococcal disease cases in China. The prevalence of macrolide resistance in these Spn strains was 99.2% (127/128). Therefore, in the current study we begin by conducting a thorough genomic analysis of genetic elements conferring macrolide resistance in these strains and aimed to identify potential hotspots and other genetic characteristics in the chromosome of these strains, that are linked to the acquisition of macrolide resistance. Subsequently, we performed a detailed investigation of cells, and metabolic determinants leading to the acquisition of macrolide resistance.

## Results

### Identification of macrolide resistance determinants and recombination in erythromycin-resistant pneumococcal strains

To identify the mechanism leading to macrolide resistance we performed whole-genome sequence analysis of 127 clinical isolates that were resistant to erythromycin (MIC≥128 µg/ml)^38^. All these isolates belonged to 20 different serotypes (S) (Fig. 1A). Pneumococcal strains belonging to S19F were more prevalent (n=43), followed by S19A strains (n=16). Our genomic analysis found that the MEGA element was carried by S19F and S19A strains that phylogenetically belong to clone complex (CC) 271. The gene *ermB* was detected in all analyzed strains. Both MEGA and *ermB* were present in *Tn916*-related elements. Consequently, we performed an analysis of recombination events within the Tn*916*-related elements and the genomic region carrying genes of the competence locus *comCDE*.

**Fig. 1.**
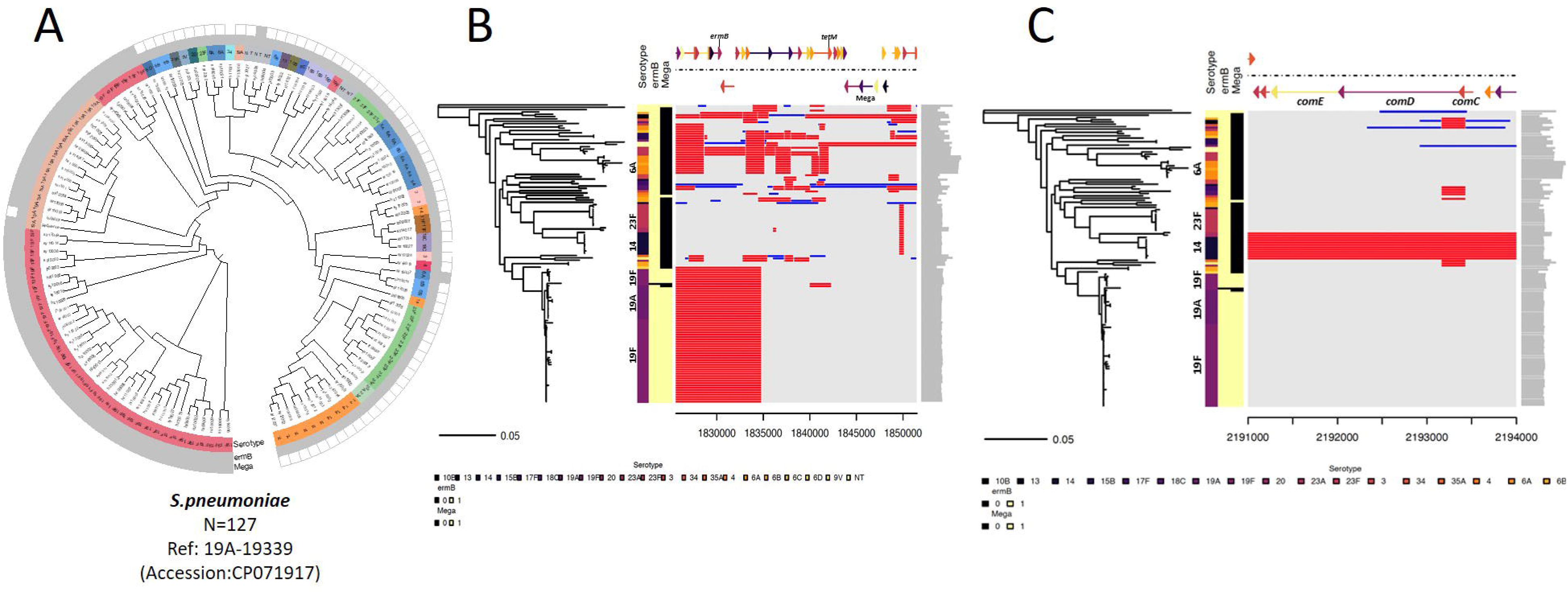
The phylogenetic tree and recombination events in macrolide-resistant clinical pneumococcal isolates. The left panel presents the phylogenetic tree of macrolide-resistant pneumococcal isolates (N=128, Accession: PRJNA795524) that was constructed in FastTree (http://www.microbesonline.org/fasttree/) using SNP calling data from Snippy (https://github.com/tseemann/snippy) (A). The metadata of serotype (different color strips) and the presence of the *ermB* gene and MEGA element (grey: positive; blank: negative) for each of the pneumococcal isolates were marked at the tip of the tree. The branches with light red shades emphasize the carrying of the MEGA element in those strains. The right panel displays the visualized recombination prediction in the region of *Tn916*-like (B) or *comCDE* (C) which were mapped against an annotated chromosome of the reference pneumococcal strain 19A-19339 (Accession: CP071917) using Snippy and Gubbins (https://github.com/nickjcroucher/gubbins), respectively. Red blocks represent the recombination blocks in each clone complex on an internal branch, which are therefore shared by multiple isolates, while blue blocks represent the recombination that occurred on terminal branches, which are unique to individual isolates. The whole data set was visualized with RCandy (https://github.com/ChrispinChaguza/RCandy) in RStudio (Version 2023.09.1+494).

The most common recombination hotspots where located in genes encoding the conjugation apparatus, including those within *ermB*, which is inserted within *orf20* carried by CC271 strains (Fig. 1B). Recombination events downstream of the T4SS genes were not identified in these Spn CC271 strains that carry MEGA. In other strains, recombination hotspots were identified in random positions of the Tn*916*-related elements including in the accessory genes’ region located downstream the genes encoding the T4SS (Fig. 1B). Most Spn strains belonging to serotype 14 and 23F did not show evidence of recombination hotspots in their *Tn916*-related elements.

We analyzed recombination hotspots in the locus *comCDE*, which encodes genes for the competence-stimulating peptide (CSP) and its cognate two-component regulatory system^39^. Surprisingly, hotspots within the *comCDE* region were only identified in serotype 14 and 23F strains, which do not carry recombination hotspots in genes within their Tn*916*-related elements (Fig. 1C). In the majority of all other strains, genes encoding the Com system were conserved with no evidence of recombination (Fig. 1B). Collectively, strains harboring Tn*916*-related elements were observed to possess recombination hotspots either within the genes encoding the conjugative machinery of Tn*916* or within the locus that regulates pneumococcal competence for DNA uptake but not in both.

### Spatial localization of pneumococcal strains within nasopharyngeal biofilm consortia

Acquisition of these Tn*916*-related elements leading to resistance to macrolides has been observed in pneumococcal strains isolated throughout the world^38, 40^, we next investigated the mechanism behind the acquisition of macrolide resistance genes. We initially studied the fitness, colonization dynamics, and ultrastructure of various pneumococcal strains, including strains carrying Tn*916*-related elements, during co-colonization in a simulated nasopharyngeal environment using an *ex-vivo* bioreactor system. This system mimics the human nasopharynx and facilitates gene transfer among strains^17, 41, 42^. Mixtures of pneumococcal strains were inoculated in the bioreactor including reference strain TIGR4, and Spn strain 8655 that carry the macrolide resistance gene *ermB* in a Tn*3872*-element (8655^Tn3872^)^18, 43^, or TIGR4 and strain D39. The growth rate of TIGR4 and 8655^Tn*3872*^, or TIGR4 and D39, when inoculated separately in Todd-Hewitt broth (THY) was similar (not shown). When strains colonized together the simulated human pharyngeal epithelium, the density of 8655^Tn*3872*^ biofilms (∼10^8^ cfu/ml) was significantly higher than that of TIGR4 (∼10^7^ cfu/ml) (Fig. 2A) while the density of D39 when co-inocualted with TIGR4 was similar (Fig. 3A). Given that a rF of ∼1×10^-3^ has been demonstrated to occur in the nasopharynx *in vivo* or *ex vivo*^15, 44^, these results suggest that vaccine and non-vaccine type pneumococci, when co-colonizing human pharyngeal cells at a bacterial density >10^6^ cfu/ml, are prone to recombination via transformation.

**Fig. 2.**
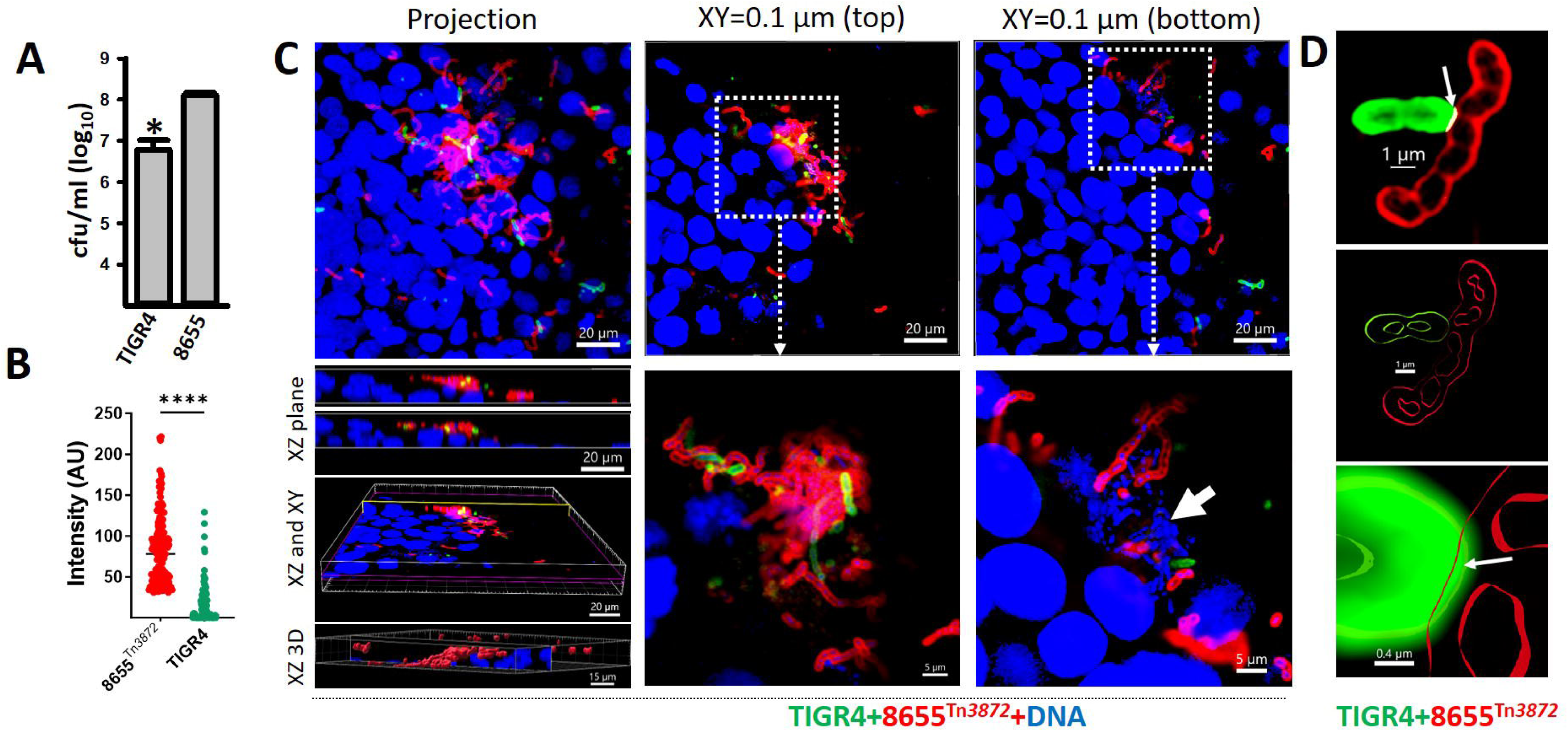
Competitive dynamics of pneumococcal strains during acquisition of resistance. (A) *S. pneumoniae* strain TIGR4 (SPJV23), and strain 8655^Tn*3872*^ (S6B) were co-inoculated into a bioreactor with human pharyngeal cells, and incubated for 6 h after which biofilms were collected, serially diluted, and plated onto different BAP with specific antibiotics to count. The error bars represent the standard error of the means calculated using data from at least three independent experiments. (B-C) Strains were stained with an anti-S4/A488 (green) and an anti-S6B/A555 (red) antibodies while the DNA was stained with DAPI (blue). Micrographs were obtained with a confocal microscope and analyzed with the Imaris software. (B) The fluorescence intensity in arbitrary units (AU) obtained from each channel was graphed. The error bars represent the standard error of the means calculated using data from two independent experiments. \*\*\*\**p* <0.0001. (C) Confocal micrographs. Left panels: the projection, the XZ plane, both Z and XY planes, and XZ plane of a 3D reconstruction are shown. Middle panle and riught penale: XY optical sections of 0.1 um each sliced from from the top and the bottom are shown. Delimited region show the are manigified in the bottom panels. The arrow point out intracelualr pneumococci. (D) Imaging analysis using the Imaris software analyzed micrographs obtained by super-resolution microscopy. Arrows point (top and bottom panel) out the area of physical colocalization.

**Fig. 3.**
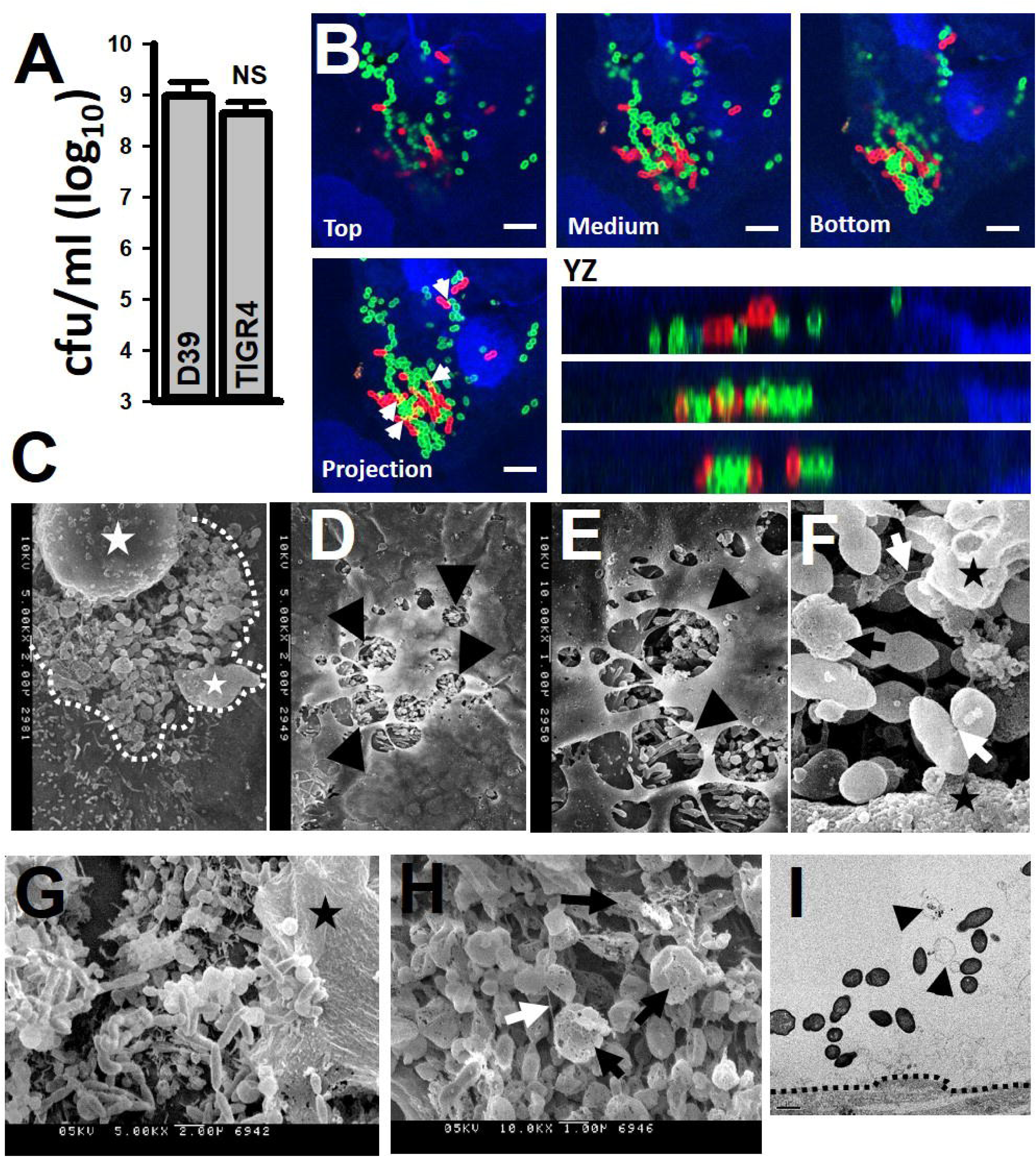
Pneumococcal biofilm consortia on human pharyngeal cells. (A) *S. pneumoniae* TIGR4 (SPJV24) and D39 (SPJV22) derivatives were co-inoculated into a bioreactor with human pharyngeal cells, and incubated for 6 h after which biofilms consortia were harvested, serially diluted, and plated onto different BAP with specific antibiotics to count. The error bars represent the standard error of the means calculated using data from at least three independent experiments. (B) Pneumococcal strains were incubated as above and then biofilms were stained with an anti-S4/A488 (green), an anti-S2/A555 (red) antibodies and the DNA was stained with DAPI (blue). Bar=5 µm. Preparations were analyzed by confocal microscopy. xy optical sections from the top, middle, or bottom sections, the xy projection, and yz optical sections, are shown. (C-I) Strains were incubated as above in the bioreactor after which pneumococcal strains were fixed and submitted to SEM (C-H) or TEM (I) image acquisition. In (C-H) white dotted line=microcolony, arrowheads=intracellular pneumococci, stars=human pharyngeal cells, black arrows=lysed pneumococci, white arrows=nanotube-like structures connecting pneumococci. In (I) black dotted line=surface of epithelial cell, arrowheads=pili-like structures.

We then tested the hypothesis that while forming a biofilm consortium, the two pneumococcal strains will be in such physical proximity to enhance the efficiency of gene transfer. To visualize individual strains within the biofilm consortium, we stained pneumococci with fluorescence-labeled serotype-specific antibodies, and the DNA with DAPI. Preparations were then analyzed with both a confocal microscope and a super-resolution confocal microscope. A biofilm consortium formed by a mixture of TIGR4 and 8655^Tn*3872*^ (Fig. 2C) or TIGR4 and D39 (Fig. 3B) was observed attached to human pharyngeal cells forming bacterial aggregates. Structural confocal analysis of the XZ plane, XZ and XY optical sections, and a XZ optical section of a 3D reconstruction showed intracellular TIGR4 and 8655^Tn*3872*^ pneumococci (Fig. 2C). Optical XY sections, each spaced 0.1 µm apart from the top and bottom of the projection, revealed that the aggregate made by TIGR4 and 8655^Tn*3872*^ locates on top but enters pharyngeal cells and these pneumococci are positioned on the same focal plane as that of cell nuclei (Fig. 2C). Remarkably, intracellular pneumococci consisted of both encapsulated and nonencapsulated bacteria (Fig. 2C, arrow). Quantification of the median fluorescence intensity, from each channel, was significantly higher for 8655^Tn*3872*^ compared with TIGR4 (Fig. 2B). The median area of the largest bacterial aggregates of 8655^Tn*3872*^ was 1684 µm^2^ while that of TIGR4 was 155 µm^2^ (not shown).

Super-resolution microscopy (Fig. 2D) and colocalization analysis shown in Fig. 3B (arrows) identified physical contact between pneumococci within biofilm consortia. Physical contact spanned over ∼1.2 µm of capsule-capsule interaction (Fig. 2D, dotted line) with 3D analysis showing the embedding of both capsules (Fig. 2D, arrow).

### Pneumococci form localized aggregates on pharyngeal cells when the acquisition of resistance via transformation occurs

We next performed ultrastructural studies using scanning electron microscopy (SEM) and transmission electron microscopy (TEM). These studies showed that pneumococci, when transformation had already occurred in the bioreactor^15^ (i.e., 6 h post-inoculation, explained in the next section), are organized in localized clusters of bacterial aggregates attached to pharyngeal cells (Fig. 3C, delimited). Pneumococci were also observed inside the cells, evidenced by disrupted membrane structures observed at various magnifications, resembling holes (Fig. 3D and 3E, arrows). Under SEM, extracellular pneumococci within clusters appear elongated (Fig. 3F-3H) and structures compatible with the type IV pili (T4P) were observed (Fig. 3F and 3H, white arrows). T4P were also evident bacterial clusters observed in micrographs collected by TEM (Fig. 3I, arrows). Within each cluster, some pneumococcal cells were observed lysed, which was characterized by the presence of bacteria with a disrupted cell wall (Fig. 3F and 3H, black arrows). Each cluster of pneumococci was surrounded by pharyngeal cells that appeared intoxicated such as those detached from the substratum or with a compromised cell membrane (Fig. 3D-G, stars).

### MEGA can be transferred in an *ex vivo* nasopharyngeal environment from disease isolates or engineered pneumococci

Infection of pharyngeal cells by pneumococci, in the bioreactor system, provides an ideal microenvironment for horizontal transfer of macrolide resistance. We therefore investigated the acquisition of the MEGA which is encoded by the *mefE* and *mel* genes^24, 25, 28^. To assess this, we inoculated in the *ex vivo* bioreactor model a pair of strains including a Spn strain isolated from pneumococcal disease carrying MEGA in the chromosome, MEGA-1.III (GA17570) or MEGA-2.II (GA41688)^25^, and a recipient D39 carrying resistance to Tet and streptomycin (D39^Str-Tet^). The recombination frequency (rF) of D39^Str-Tet^ to acquire MEGA from GA17570, or from GA41688, was 2.44×10^-7^ or 1.18×10^-5^ respectively (Fig. 4A, and 4B). The colonization density (cfu/ml) of the recipient D39^tr-Tet^ when co-inoculated with either donor strain was similar thereby the differences cannot be attributed to a decreased population of the recipient (Figs. 4A and 4B, right panels). Tetracycline resistance was acquired by MEGA-carrying strains (i.e., from D39^Str-Tet^) at a rF of 7.59y×10^-5^ (GA17545, Fig. 4A), or 8.15×10^-5^ (GA41688, Fig. 4B).

**Fig. 4.**
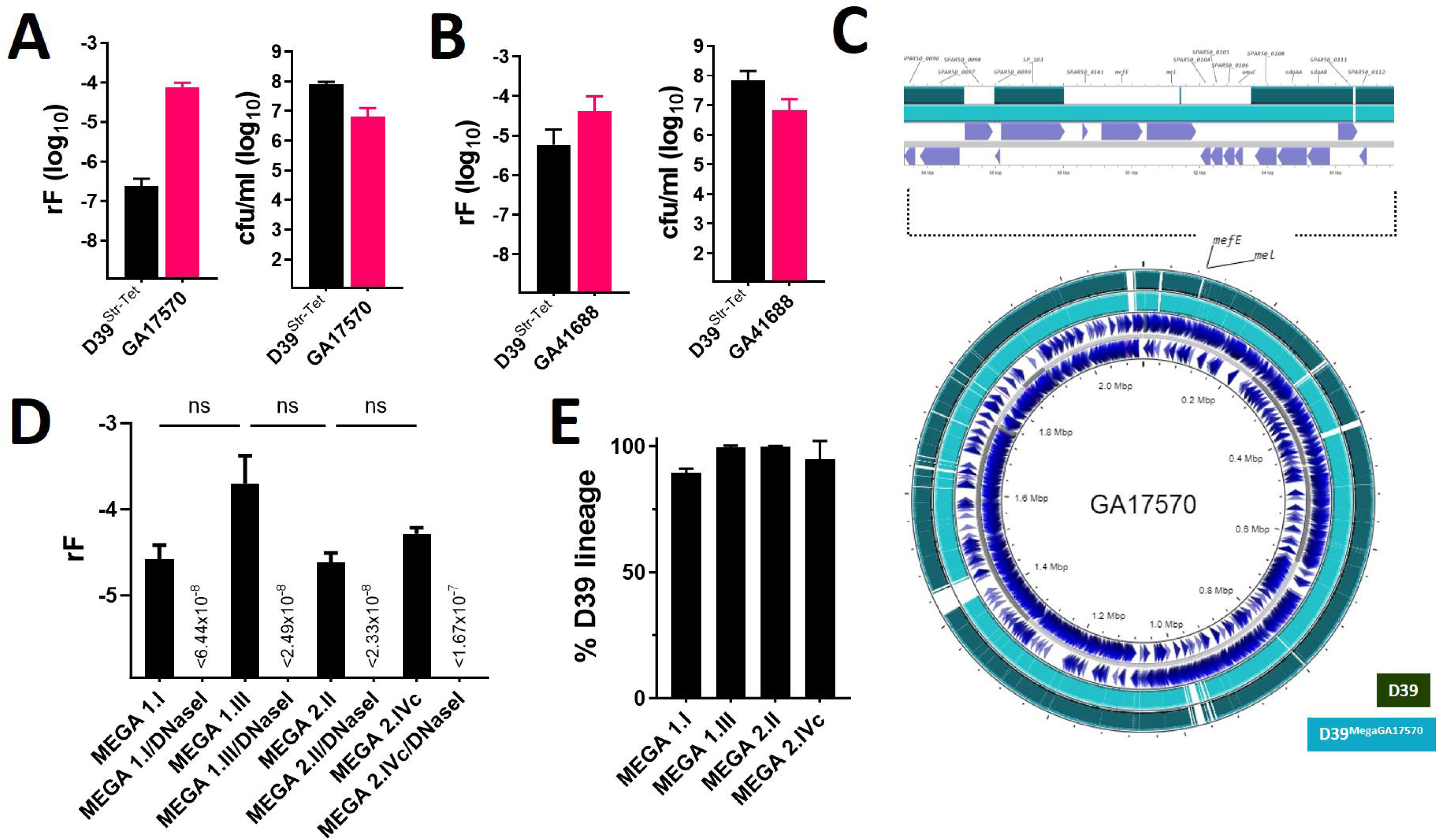
Transference of macrolide resistance carried in MEGA. (A) *S. pneumoniae* D39 (SPJV54) and GA17570, or (B) SPJV54 and GA41688 were co-inoculated into the bioreactor and incubated for 6 h after which biofilms were collected, serially diluted, and plated onto blood agar plates containing the appropriate antibiotic mixture to count each parent strain, or transformants from each lineage. The recombination frequency (rF) was calculated (left panels) and density of each strain in also shown (right panels). (C) Recipient strain D39 and transformant D39^Mega17570^ were mapped against donor strain GA17570. Insertion of MEGA (*mefE*/*mel*) into D39^Mega17570^ and a fragment of at least ∼9 kb from donor are shown. (D) D39 (SPJV22) was inoculated in the bioreactor along with TIGR4 carrying MEGA 1.I, MEGA 1.III, MEGA 2.II, or MEGA 2.IVc, and incubated for 6 h. In parallel experiments performed with each pair of strains, the culture medium was supplemented with DNase I. Biofilms were harvested and the rF was calculated as above. (E) SPJV22 recombinants (i.e., carrying MEGA) were pooled, the DNA was extracted and used in serotype specific qPCR reactions to detect the total copies of serotype 2 (SPJV22) or serotype 4 (TIGR4-MEGA derivative). The percentage of SPJV22 (D39 lineage) was used to construct the graph. In panels (A, B, D and E) the error bars represent the standard error of the means calculated using data from at least three independent experiments.

Whole genome sequencing confirmed that in strain D39^Mega/GA17570^ the MEGA element was located in the same chromosome region as that in the donor GA17570, in a region between the capsular polysaccharide biosynthesis protein *capD* gene (SPD_0099, D39 nomenclature) and SPD_1001, a gene encoding for a putative hydrolase (Fig. 4C).

To investigate if the location of MEGA in the chromosome influences the rF, we assessed the transfer of MEGA using strains with the same genetic background. To this purpose, TIGR4 strains were engineered to carry MEGA class 1.I, 1.III, 2.II, or 2.IVc in the chromosome (Table 1). A recipient D39^Str-Tet^ was incubated in the bioreactor with a TIGR4^MEGA^ strain and the rF was investigated. D39^Str-Tet^ recombinants carrying now macrolide resistance in a MEGA element (D39^Str-Tet/MEGA^) were obtained at a similar rF >2.01×10^-3^ (Fig. 4D). Using a qPCR approach, we confirmed that >90% of D39^Str-Tet/MEGA^ recombinant strains carried capsules genes for serotype 2 (Fig. 4E). Acquisition of MEGA by D39^Str-Tet^ was inhibited by treatment with DNAseI indicating that this genetic element (≥5.4 kb) conferring macrolide resistance was acquired by transformation (Fig. 4D). Taken together, the acquisition of MEGA by a recipient pneumococcal strain is not influenced by the chromosomal region where this genetic element is located.

**Table 1.**
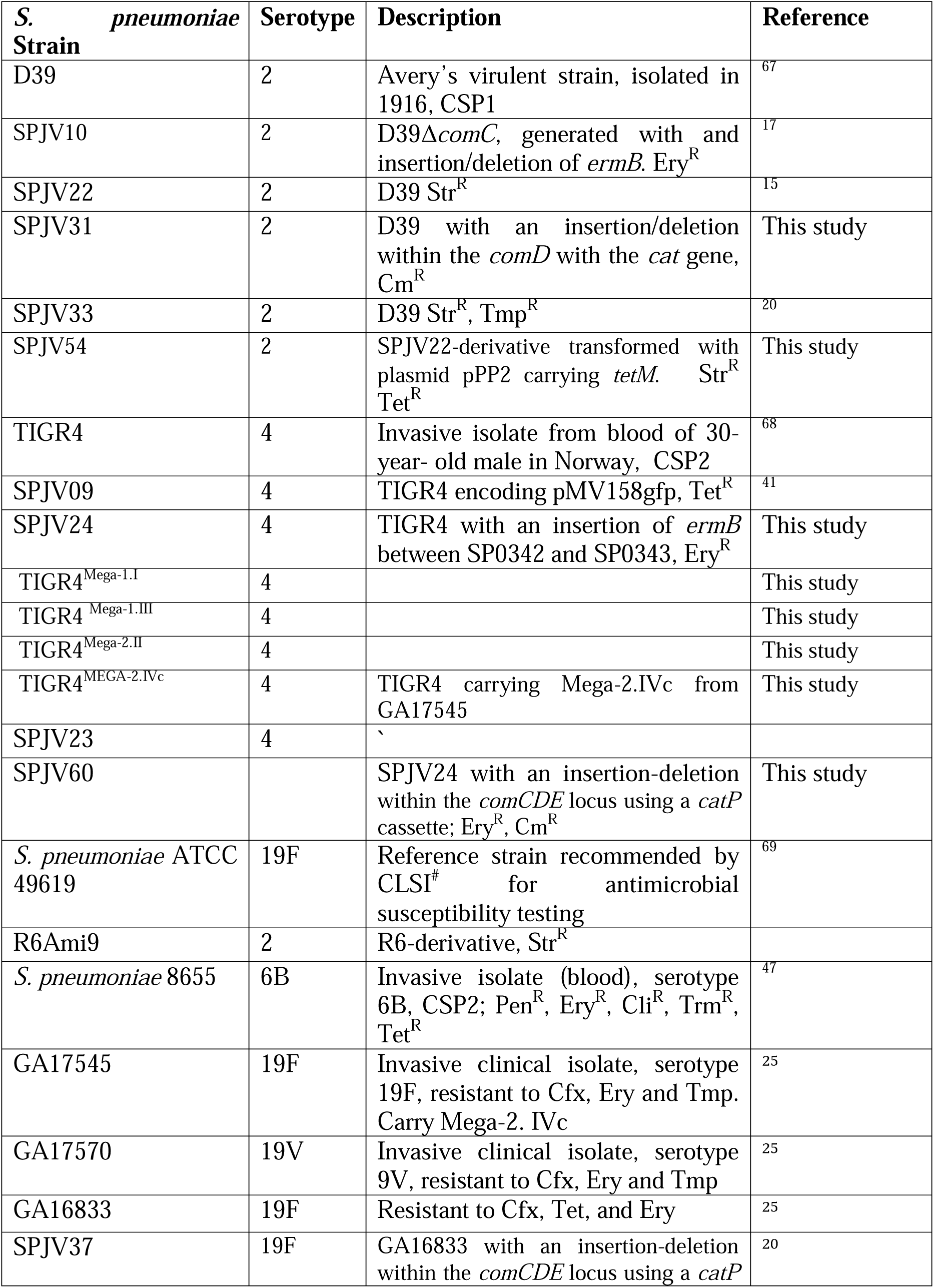
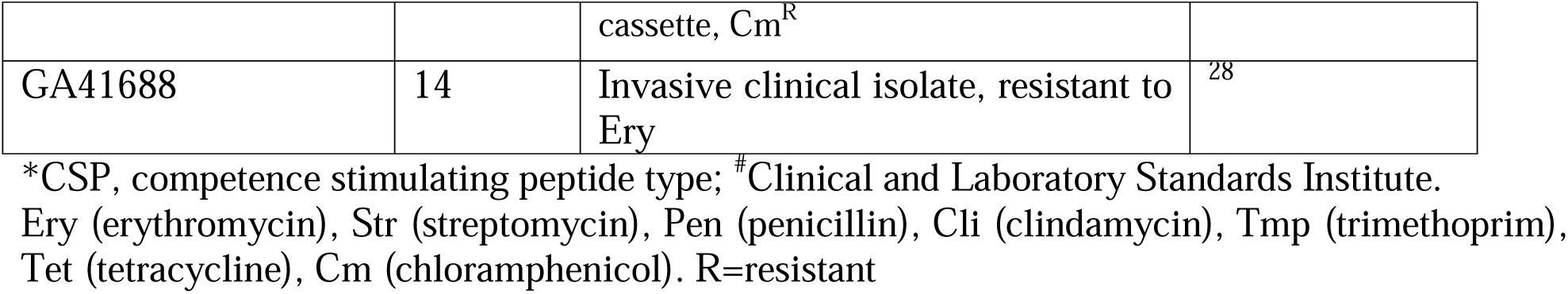
*Streptococcus pneumoniae* strains utilized in this study.

### Similar recombination frequency for the acquisition of macrolide resistance encoded in the chromosome or in large ICEs

We have demonstrated that acquisition of large ICEs by *S. pneumoniae* occurs in the bioreactor and that it is facilitated by the transformation machinery^19,20^. We therefore sought to compare, under the same culture conditions of the bioreactor, the rF for the acquisition of macrolide resistance encoded in genetic elements of varying sizes (∼1, ∼5.5, ∼23.5, or ∼51 kb). To this end, we engineered, or selected, pneumococcal strains carrying macrolide resistance genes *ermB* (TIGR4^Ω*ermB*^) or *mef(E)/mel* (i.e., MEGA) either inserted in the chromosome, (TIGR4^Mega1.I^ or TIGR4^Mega2.IVc^) or carried within transposons Tn*2009* (GA16833) (Fig. 5A) or Tn*Meg* (GA17545). Since Tn*2009* carries macrolide resistance and the *tetM* for tetracycline resistance, we utilized a recipient strain D39 carrying mutations ∼800 bp apart in the *rpsL* and *folA* gene, conferring resistance to streptomycin and trimethoprim (D39^Str-Tmp^), respectively. Mutations and genes associated with resistance were confirmed in the recipient and donor strains by whole genome sequencing^19^.

**Fig. 5.**
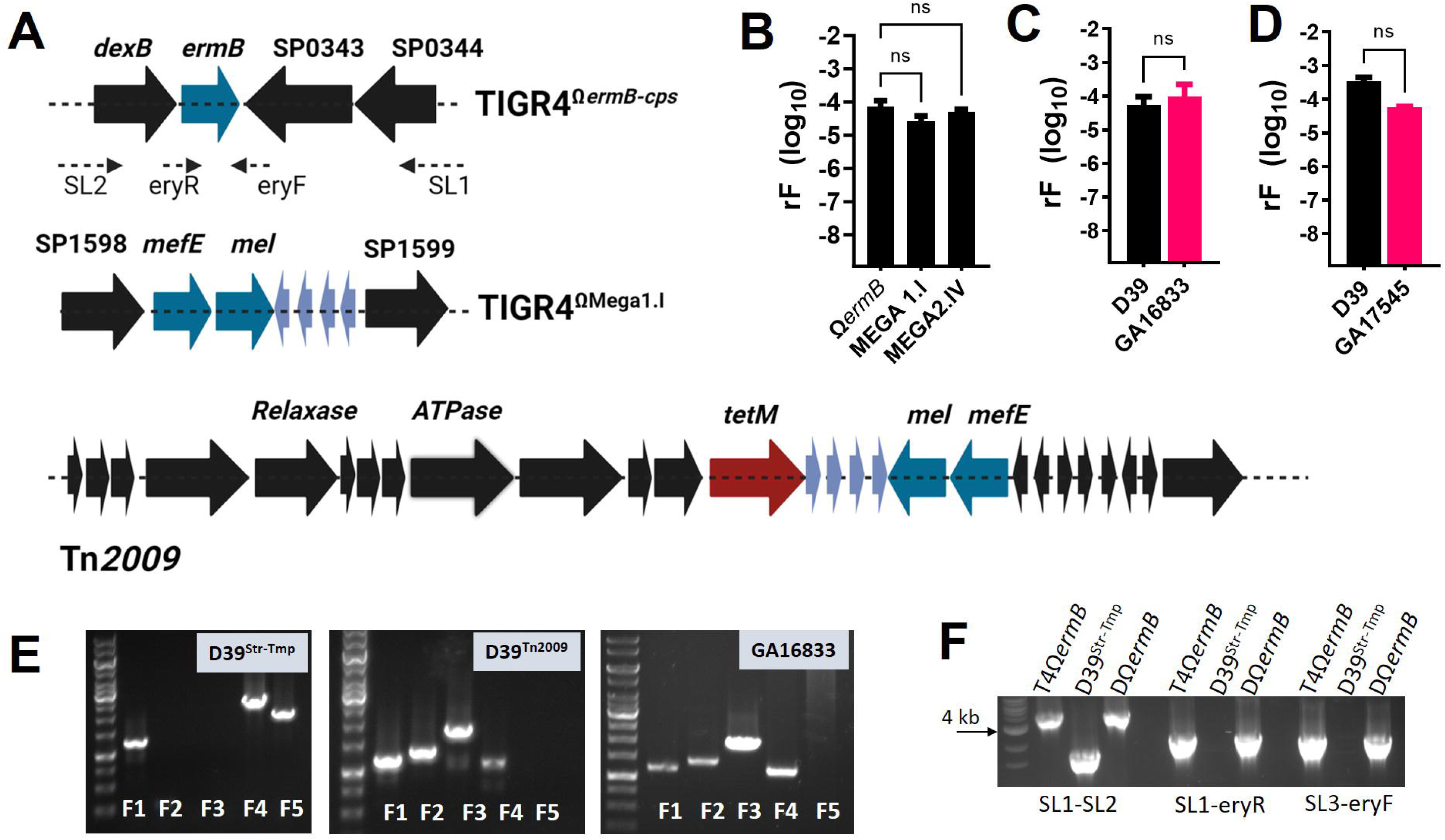
Transference of macrolide and tetracycline resistance carrying transposon Tn2009. (A) Genomic context of the insertion of *ermB* in TIGR4^Ω*ermB*^, or MEGA (*mefE*/*mel*) in TIGR4^ΩMega1.I^. Primers SL2, eryR, eryF and SL1, underneath the top panel were utilized below. The map of Tn2009 from GA16833 is shown, indicating the location of genes encoding for the relaxase, and ATPase, of the putative type IV secretion apparatus, and genes *tetM*, *mefE* and *mel*. (B-D) D39 (SPJV33) was inoculated in the bioreactor along with (B) TIGR4^Ω*ermB*^, TIGR4^ΩMega1.I^, or TIGR4^ΩMega2.IV^ or (C) GA16833, or (D) GA17455. Experiments were incubated for 6 h and biofilm bacteria were harvested from the bioreactor, diluted and platted onto blood agar plates containing the appropriate antibiotics to investigate the recombination frequency (rF). The error bars represent the standard error of the means calculated using data from at least three independent experiments. NS=no significant. (E) DNA was purified from SPJV22 (D39^Str-Tmp^), D39^Tn2009^, or GA16833, and utilized as template in PCR reactions with pair of primers F1-F5. (F) DNA was purified from TIGR4^Ω*ermB*^ (T4Ω*ermB*), SPJV33 (D39^Str-Tmp^) or SPJV33^Ω*ermB*^ (DΩ*ermB*) and used as template in PCR reactions with primer pairs listed below.

The rF of recipient D39^Str-Tmp^ acquiring macrolide resistance from the engineered TIGR4 donor strains and that carried either a ∼1 kb (Ω*ermB*), ∼5.4 (MEGA1.I), or ∼5.5 kb (MEGA2.IVc) occurred at a similar rF with a median of 5.03×10^-5^ (Fig. 5B). The donor strain, TIGR4^Ω*ermB*^, TIGR4^Mega1.I^ or TIGR4^Mega2.IVc^ did not acquire streptomycin or trimethoprim resistance (not shown) from recipient D39^Str-Tmp^ under the bioreactor incubation condition^15^.

Remarkably, the rF for the acquisition of MEGA carried in Tn*2009* (∼23.5 kb) or Tn*Meg* (∼53 kb), from donor strain GA16833, or GA17545, respectively, was similar to that yielded by engineered TIGR4 strains, with a median rF of 5.47×10^-5^, or rF=3.50 ×10^-4^, respectively (Fig. 5C and 5D). Unlike TIGR4 that did not acquire resistance from the recipient strain, donor strains GA16833 (Tn*2009*) and GA17545 (Tn*Meg*), acquired streptomycin resistance from D39^Str-Tmp^ at a rF of 9.65×10^-5^ or 5.89×10^-5^, respectively, thereby confirming the micro-environment of the bioreactor allowed for bidirectional exchange of DNA.

We then conducted PCR analysis to confirm that *ermB*, or the MEGA element, was acquired by the recipient. PCR analysis of ten different transformants confirmed that in D39^Ω*ermB*^ the *ermB* gene was located downstream *dexB* in the same location as that of the donor strain TIGR4^Ω*ermB*^ but absent in the recipient D39^Str-Tmp^ (Fig. 5F). For example, PCR using primers SL1-SL2 amplified a ∼4.3 kb fragment in both TIGR4^Ω*ermB*^ and D39^Ω*ermB*^ but a ∼2 kb fragment in D39^Str-Tmp^ due to the absence of *ermB* and SP0343 (Fig. 5A and 5F). The MEGA element, *mef(E)/mel,* was also amplified from ten D39^Mega1.I^ transformants and PCR analysis located the insertion within the same region of the chromosome (not shown).

PCR reactions amplified similar PCR products using DNA from both GA16833 and D39^Tn*2009*^ that were absent when the DNA template was purified from D39^Str-Tmp^ (Fig. 5E, listed as F2, F3, and F4). Two PCR products (F1 and F5), amplified using DNA from D39^Str-Tmp^ were absent in GA16833 and D39^Tn*2009*^ (Fig. 5E). These results revealed that the acquisition of macrolide resistance occurs at a similar frequency in strain D39 whether MEGA and/or *ermB* are carried within an ICE or in the chromosome, and regardless of the molecular size of the ICE.

### Metabolomics studies during the acquisition of pneumococcal ICEs

To gain further insight into the mechanism by which macrolide-resistant carrying ICEs are acquired by Spn, we conducted metabolomics studies. Supernatants from bioreactors that had been left uninfected, infected with D39 and GA16833 or infected with D39Δ*comD* (SPJV31) and GA16833 were harvested and analyzed. Supernatants were collected for 60 min, between 1 and 2 h post-inoculation (labeled as 2 h), or 3 and 4 h post-inoculation (labeled as 4 h). First, partial least squares discriminant analysis (PLSDA) was conducted to investigate if the groups were separable. After merging the two time points of each infection condition, the overall error rate using centroid distance demonstrated that metabolites in cells infected with D39 and GA16833 were more different than the other two groups (Fig. 6A). PLSDA further identified metabolites from pharyngeal cells incubated with D39 and GA16833 for 2 h as the most distinct (Fig. 6B).

**Fig. 6.**
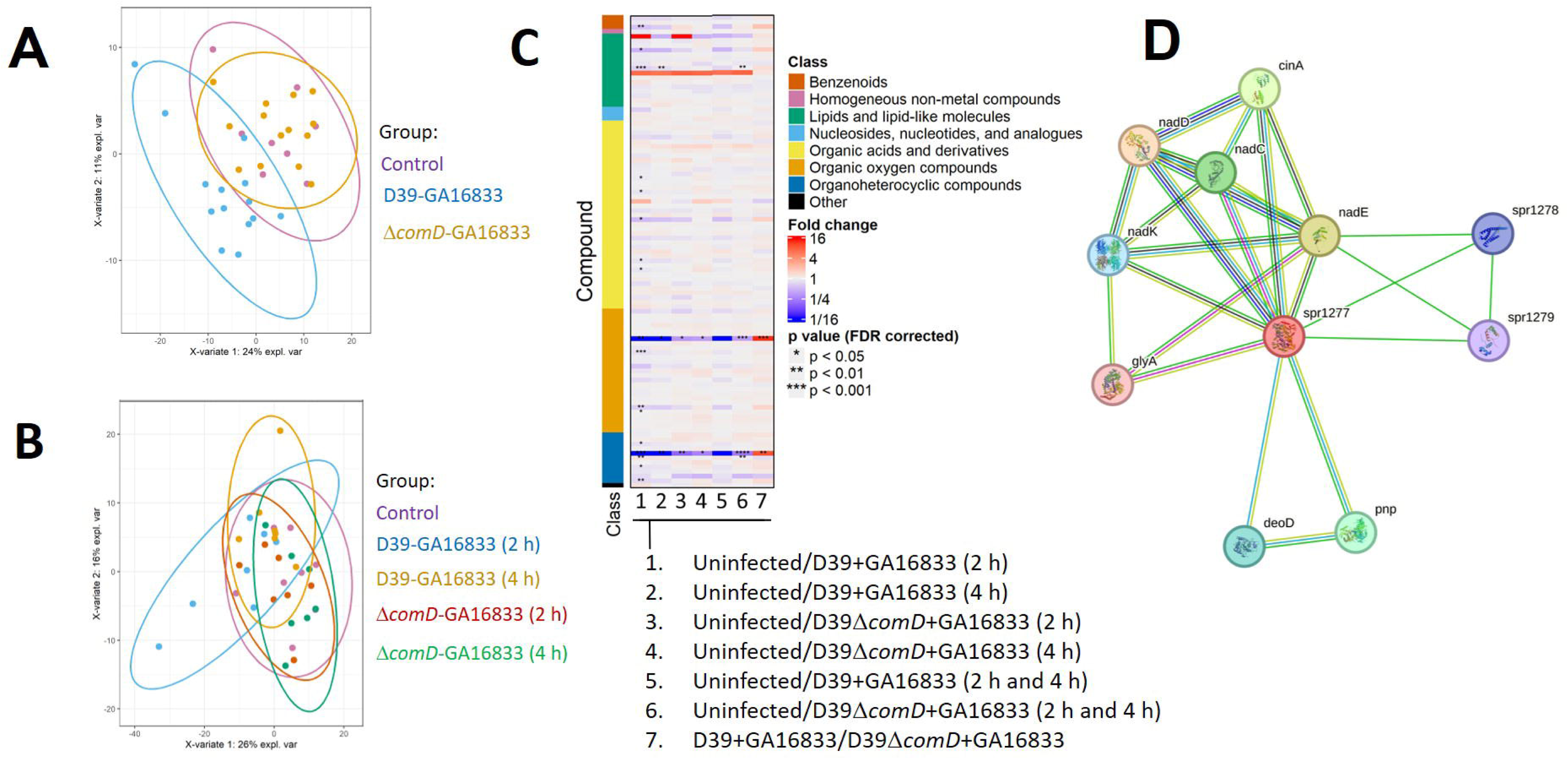
Primary metabolism analysis of molecules from pneumococci and pharyngeal cells during the acquisition of a pneumococcal ICE. The bioreactor was inoculated with D39 (SPJV22) and GA16833 or D39Δ*comD* and GA16833, and incubated at 37°C. Supernatants coming off the bioreactor chamber were collected for 60 min between 1 and 2 h, or 3 and 4 h, post inoculation. Primary metabolism analysis was performed to sterilized supernatants. (A) PCA biplot of the first two components extracted from a principal components analysis; group membership is indicted, and normal data ellipses for each group. (B) Biplots of the first two components extracted from a partial least squares discriminant analysis, group membership is indicated, and normal data ellipses for each group is shown. (C) Heatmap for the identified compounds. Each column is a pairwise comparison between two groups. Each row is a chemical compound. Asterisks indicates statistical significance from Mann Whitney U test. The color bar on the side indicates the chemical superclass from the class column. (D) STRING protein interaction network analysis of Nicotinate phosphoribosyltransferase (Spr1733). Shown are 25 edges (interactions) linking 11 proteins (nodes). The average local clustering coefficient was 0.858 and protein-protein interaction enrichment *p* value was 2.24×10^-4^ demonstrating a potential biological interaction among proteins.

When the relative abundance of the metabolites was assessed, those from cells incubated with D39 and GA16833 for 2 h exhibited the most significant differences compared with metabolites in uninfected cells (Fig. 6C and Table 2). Three known metabolites and nine unknow molecules were differentially identified (Table 2). Whether infected with D39 and GA16833, or with D39Δ*comD* and GA16833, and incubated either 2 h or 4 h, nicotinic acid and sucrose were significantly reduced compared with uninfected cells incubated in the bioreactor under the same culture conditions (Table 2). We conducted a comparison between the metabolites found in the supernatants from pharyngeal cells infected with D39 and GA16833 and those from cells infected with D39Δ*comD* and GA16833. Nine known molecules including nicotinic acid (∼8-fold increase) and sucrose (∼11-fold increase) were enriched in supernatants from cells infected with D39Δ*comD* and GA16833. Together, metabolomics studies indicate that the transformation capacity of the recipient, strain D39, causes a significant alteration of metabolites involved in nucleic acid synthesis and sucrose metabolism (Table 2).

Nicotinic acid (vitamin B complex or niacin) is a precursor of NAD (Nicotinamide Adenine Dinucleotide) and NADP, two molecules important for bacterial physiology. Since the role of nicotinic acid in the acquisition of resistance by Spn is unknown, we conducted a protein interaction network analysis using STRING. A conserved hypothetical protein, nicotinate phosphoribosyltransferase (Spr1277, R6 nomenclature)^45^ that catalyzes the first step in the biosynthesis of NAD from nicotinic acid yielded 25 edges and 11 nodes in the STRING database. The proteins shown in Fig. 6D are biologically connected. As expected, Spr1277 showed direct interaction with NAD biogenesis proteins such as NanC, NadD, NadE, and NadK. Besides these proteins, a direct STRING interaction was obtained with the competence induced protein A (CinA). No interactions were observed in STRNG when sucrose-6-phosphate hydrolase (ScrB), which enables bacteria to metabolize sucrose, was used to analyze the network against all other enzymes derived from the metabolomics analysis, i.e., Spr1277, NanC, or CinA.

### Enhanced transformation frequency for the acquisition of pneumococcal transposon compared to acquisition of the capsule locus

The similar rF obtained in Fig. 5 experiments prompted us to further investigate if the entire ICE was acquired by most macrolide-resistant recombinants. Because the acquisition of pneumococcal transposon is facilitated by the transformation machinery^19^, we compared the rF of the large Tn*Meg*, ∼52 kb (Supplemental information 1) against that for the acquisition of the capsule *cps* locus (∼23 kb), whose genes are acquired via transformation. To assess this, we took advantage that TIGR4^Ω*ermB*^ had been engineered to carry *ermB* ∼5 kb upstream the first gene of the *cps* locus, *cps4A*. Given that there is high homology in the *cps* locus of the donor TIGR4^Ω*ermB*^ and the recipient strain D39, but Tn*Meg* carries ∼52 kb of heterologous DNA (Fig. 7A), we would have expected a higher rF for the acquisition of the *cps* locus compared with that of the acquisition of the full-length Tn*Meg*.

**Fig. 7.**
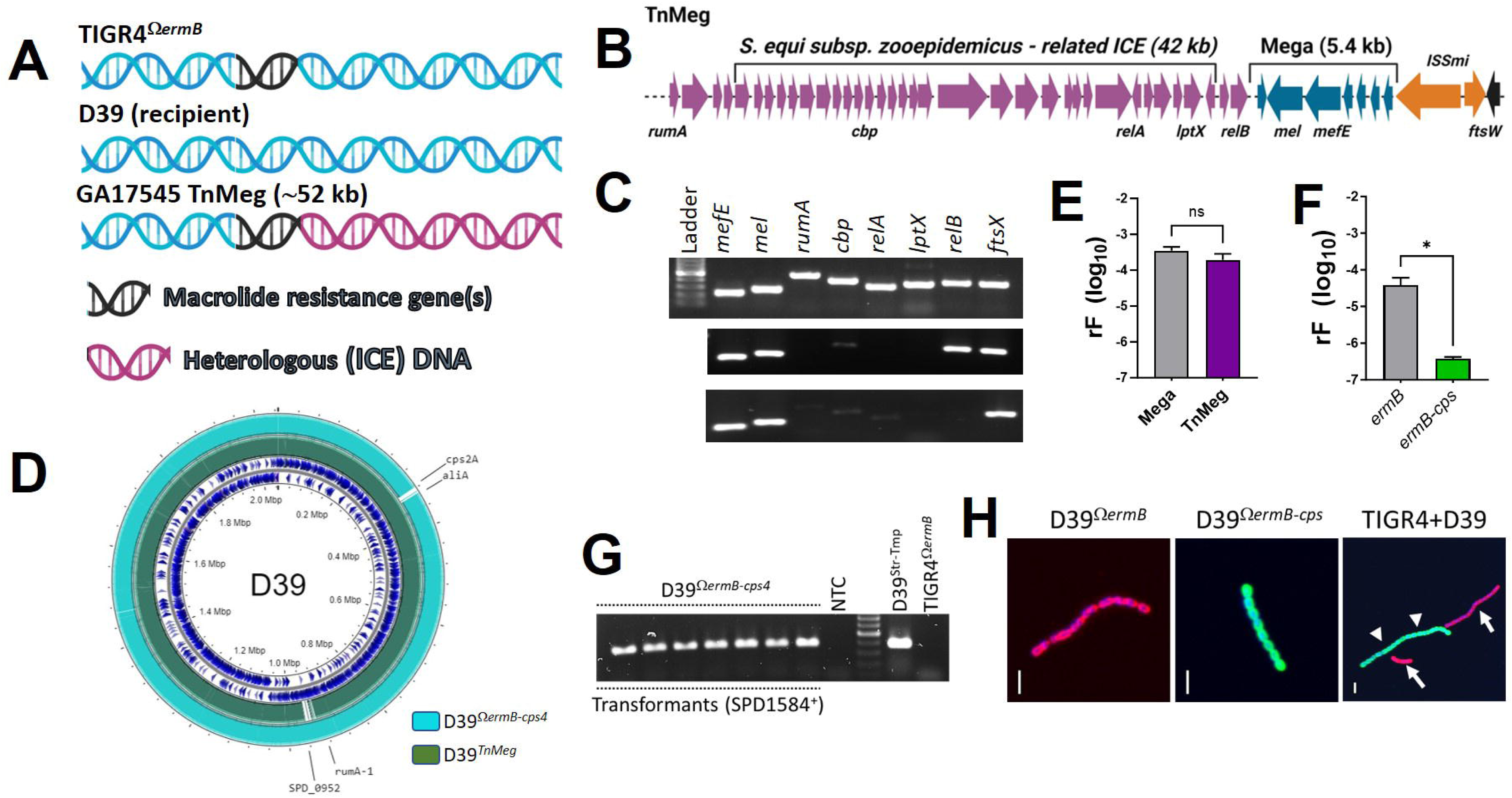
Recombination frequency for the acquisition of Tn*Meg* or the capsule locus by recipient D39. (A) Diagram representing homologs (blue) and heterologous (red) DNA regions where macrolide resistance genes (black), *ermB* or *mefE*/*mel* are carried in donor strains TIGR4^Ω*ermB*^, or GA17545, as compared to the recipient D39. (B) Schematic map of Tn*Meg* from GA17545. Genes utilized as targets in PCR are shown underneath. (C) DNA was extracted from transformants D39^Tn*Meg*^ and utilized as template in PCR reactions amplifying the genes shown. (D) Mapping of D39^Tn*Meg*^ and D39 ^Ω*ermB*-*cps4*^ against recipient D39. (E) D39 (SPJV33) and GA17545 were inoculated in the bioreactor and incubated for 6 h. Biofilm bacteria were harvested from the bioreactor, diluted and platted onto blood agar plates (BAP) containing the appropriate antibiotics to investigate the recombination frequency (rF). (E) The rF of all transformants (D39^Tn*Meg*^) growing on BAP containing streptomycin, trimethoprim and erythromycin is shown as (MEGA). Those D39^Tn*Meg*^ carrying the full-length MEGA, as confirmed by PCR, were utilized to adjust the rF and that is shown as (TnMeg). (F) D39 (SPJV33) and TIGR4^Ω*ermB*^ were inoculated in the bioreactor and incubated for 6 h. Biofilm bacteria were harvested from the bioreactor, diluted and platted onto blood agar plates (BAP) containing the appropriate antibiotics to investigate the recombination frequency (rF). The rF of all transformants (D39^Ω*ermB*^) growing on BAP containing streptomycin, trimethoprim and erythromycin is shown as (*ermB*). Those D39 ^Ω*ermB*^ carrying the capsule locus from TIGR4, were utilized to adjust the rF and that is shown as (*ermB-cps*). (G) D39 ^Ω*ermB*-*cps4*^ transformants were DNA extracted and this DNA was used as a template in PCR reactions amplifying the D39 gene SPD_1584, encoding a putative ABC transporter, permease protein. (H) D39^Ω*ermB*^, D39 ^Ω*ermB*-^ *^cps4^* or a mixture of D39 and TIGR4^Ω*ermB*^ was stained with an anti-S2 antibody labeled with Alexa555, and an anti-S4 antibody labeled with Alexa488. Preparations were analyzed by confocal microscopy. Arrows= D39, arrowheads= TIGR4^Ω*ermB*^. In all panels Bar=3 µm.

To identify transformants that had acquired the full-length Tn*Meg* (∼52 kb), we first extracted DNA from D39^Tn*Meg*^ transformants (N=100), and we performed eight PCR reactions with primers spanning Tn*Meg* (Fig. 7B). Figure 5C shows representative PCR reactions of D39^Tn*Meg*^ acquiring full-length Tn*Meg* or those that only acquired MEGA and short pieces of DNA either or both upstream and downstream. Whole genome sequencing confirmed the acquisition of Tn*Meg* by D39 (Fig. 7D). Notably, 54% of D39^Tn*Meg*^ transformants acquired the ∼52 kb Tn*Meg*. Having obtained the percentage of D39^Tn*Meg*^ carrying full-length Tn*Meg*, we re-calculated the rF to now reflect true acquisition of the entire ICE and the adjusted rF was 1.95×10^-4^ (Fig. 7E). There was, however, no statistical significance when we compared the rF for the acquisition of MEGA (i.e., transformants growing on erythromycin plates) compared with that for acquiring the full-length Tn*Meg*.

We preformed experiments in parallel with donor TIGR4^Ω*ermB*^ and recipient D39^Str-Tmp^ and obtained a rF=3.84×10^-5^ (Fig. 7F). Because the *cps* locus is highly similar between TIGR4 and D39 and conventional PCR mapping is not possible, we utilized qPCR assays that differentiate serotype 2 from serotype 4^46,47^ thereby D39^Ω*ermB*^ transformants with a *cps4* positive qPCR reaction represented D39 acquiring capsule genes from TIGR4. Only 12% of these colonies (8/66) yielded a qPCR positive reaction for serotype 4. Conventional PCR using D39-specific primers (Fig. 7G) and wgs (Fig. 7D) confirmed that transformants were D39^Ω*ermB*^ that carried the *cps* from TIGR4 (D39^Ω*ermB-cps4*^). A mixture of anti-capsule antibodies against serotype 2 or serotype 4 capsule (anti-S2A555 and anti-S4A488) demonstrated that all eight D39^Ω*ermB-cps*^ expressed serotype 4 capsule whereas D39^Ω*ermB*^ strains with a negative PCR reaction for *cps4* expressed serotype 2 capsule (Fig. 7H). As a control, a mixture of donor and recipient showed that antibodies are specific for each capsular type (Fig. 7H). The adjusted rF for the acquisition of the *ermB-cps4* capsule locus (rF=3.77×10^-7^) was significantly different to that of the acquisition of *ermB* (rF=3.84×10^-5^) (Fig. 7F). We then deleted the *comCDE* locus in TIGR4^Ω*ermB*^ and we confirmed that TIGR4Δ*comCDE*^Ω*ermB*^ had a significant transformation defect compared with the parent strain (not shown). This new TIGR4Δ*comCDE*^Ω*ermB*^ capsule donor strain was incubated along with the recipient D39^Str-Tmp^ in the bioreactor and transformants with the genotype D39^Str-Tmp/Ery^ were harvested. The adjusted rF for the acquisition of the capsule locus from TIGR4Δ*comCDE*^Ω*ermB*^ into strain D39^Str-Tmp^ was again significantly different than that of the acquisition of *ermB* (not shown).

Taken together, we demonstrated that the rF for acquiring TnMeg (∼52 kb) by recipient D39^Str-Tmp^ was three orders of magnitude higher than that to acquire a homologous *ermB-cps4* locus indicating that the mechanism of acquisition of this large ICE is further facilitated by other components outside the transformation machinery.

## Discussion

We demonstrated in the current study structural, metabolic and genetic characteristics occurring during the acquisition of macrolide resistance by pneumococcal strains. Spn strains form highly dynamic nasopharyngeal biofilm consortia, with pneumococci fused into “islets” of ∼20 µm while maintaining their own identity (i.e., capsule expression). Importantly, the close proximity within the pneumococcal islets facilitated rapid acquisition of antibiotic resistance carried in an ICE or chromosomally encoded within the MEGA element. Macrolide resistance elements, named MEGA or *ermB*, were transferred at a high frequency in the bioreactor without selective pressure. Moreover, supplementing the bioreactor with sub-MIC erythromycin did not significantly alter the rF for the acquisition of Tn*2009* by Spn (not shown). Plasmid transfer frequency of the conjugation plasmid encoding the tetracycline-efflux pump TetA, in *Escherichia coli*, remains unchanged despite the presence of tetracycline^48, 49^. In contrast, the presence of cell wall targeting antibiotics affected the transformation efficiency of the SCC*mec* cassette, encoding methicillin resistance in *Staphylococcus aureus*^50^. Overall, this evidence has important implications for our understanding of the mechanism of antibiotic resistance dissemination among pneumococci, and perhaps other streptococci.

Acquisition of genes in the nasopharyngeal microenvironment by recombination via transformation has driven the spread of drug resistance, and the acquisition of capsule genes from pneumococcal vaccine and vaccine-escape strains^51, 52^. Current methods for studying the rF for the acquisition of large genetic elements carrying antibiotic resistant genes, in most cases, skew the analysis of frequencies toward the antibiotic selection utilized. Our study included PCR mapping of >200 transformants that acquired macrolide resistance carried either upstream the capsule locus (∼28 kb), or in a large ICE (∼53 kb). To our surprise, the adjusted rF for the acquisition of the full-length ICE was three orders of magnitude higher than that to acquire the capsule locus. Accordingly, our molecular studies revealed that >60% of D39^Tn*Meg*^ transformants acquired the full-length Tn*Meg*, whereas <20% transformants screened acquired the full-length capsule locus. Elegant *in vitro* transformation studies demonstrated that the single largest recombination event in Spn resulted in the acquisition of 30 kb^53^. Therefore, acquisition of the entire Tn*Meg* at such different rF, compared with the acquisition of transformation-driven capsule locus, may suggest that additional elements are involved. Recent evidence showing that the acquisition of a ∼60 kb SCC*mec* cassette driving the spread of methicillin resistance in *S. aureus* supports the above hypothesis. Acquisition of SCC*mec* was shown to occurs through transformation but it required additional elements including the CcrAB-mediated excision/integration system encoded within SCC*mec*^50^.

We recently provided evidence that the transformation machinery facilitates the transfer of large ICEs in Spn^19^. Spontaneous natural competence develops in the bioreactor at a high frequency (∼10^-4^) that all elements necessary for the acquisition of ICEs occurred. The absence of human pharyngeal cells, for instance, causes a reduced rF of ∼10^-7^ (not shown and^15^). Thus, an orchestrated machinery driven the spread of macrolide resistance involves both bacterial and host cell factors. We have at least two hypotheses to explain why the competence system facilitates these acquisition events. In the first one, we suggest that the competence regulon affects the formation of the DNA exchange islets on human cells, or the proximity of pneumococci within the biofilms. In support to this, competence development has been associated to early events during the attachment of Spn to host cells^17, 54^. Whereas the density of *com* mutants was similar as that of the wt strains in dual strain biofilms^19^, early events required for the acquisition ICEs may have been perturbed. A second hypothesis relates to the metabolite host response against a functional competence system, and therefore to the absence of a metabolite(s) or the absence of metabolic routes to process an essential molecule(s), required to trigger the acquisition of an ICE. We are currently assessing these hypotheses in our laboratories.

Recent ultrastructural studies have shown that the Gram-positive bacterium *Bacillus subtilis* produces membranous nanotubes, enabling the exchange of molecules among bacterial cells^55^. The ultrastructure of pneumococci at the time of the exchange of genetic material have not been investigated, in part because of the lack of a nature-like model. We demonstrated here that pneumococci produce abundant TIV pili at the time of the acquisition of resistance and some vesicle-like structures were observed. The fact that treatment with DNaseI inhibited the acquisition of chromosomally-encoded MEGA and ICE, in the bioreactor, suggest that nanotubes or membrane vesicles are not required to transfer macrolide resistance.

A challenge to recreate persistence nasopharyngeal colonization is the irreversible pneumococcal autolysis triggered at 8 h post-inoculation of biofilms in a microplate model ^17, 56^. We have used a circulated bioreactor system with human nasopharyngeal cell monolayer to mimic the *in vivo* nasopharynx environment, in which pneumococcal strain would colonize human cells without autolysis. In the simulated pneumococcus natural niche, resistance to tetracycline, streptomycin, erythromycin, trimethoprim, or beta lactams such as ampicillin and cefuroxime (not shown) was transferred very rapidly, in less than 8 h of co-culture. Whereas we demonstrated that the acquisition was mediated by transformation, as we would expect, transformation did not occur when strains were incubated together for 8 h in micro-titer plates whether or not pharyngeal cells were seeded on those plates. We therefore proposed that in the absence of autolysis human pharyngeal cells trigger pneumococcal transformation leading to the stochastic acquisition of resistances at a high rF of ∼10^4^.

Moreover, recombination via transformation or mosaic acquisitions that can affect capsular expression, including serotype switching between pneumococcal strains ^57^. For example, Yang Baek, et al. (2018) reported a capsule switching from S11A to S15A via recombination in an extensively drug-resistant *S. pneumoniae* strain ^58^. Our whole genome sequencing data from macrolide resistant clinical isolates showed that high frequency recombination occurred around Tn*916*-like elements and capsule expression region, while no serotype switch event was detected in all tested strains. It was interested us whether the transference of antibiotic resistance genes occur independently or along with *cps* genes transformation. Therefore, we inserted an *ermB* gene near the capsule locus of TIGR4 strain for transformation assays on human nasopharyngeal cells. Our results revealed that the majority of D39 transformants (75%), carried only pieces of the capsule locus from TIGR4, and not the entire capsule locus. We located some of these DNA fragments in an intergenic region between the *clpL* and *mraY* genes and ending within the *csp2A* gene without affecting the capsule expression of recombinants (not shown). This may explain the observation we made in our clinical strains indicating that the cps locus is a hotspot for recombination but serotype switch was not detected. Our studies conducted after the introduction of the pneumococcal vaccine demonstrated a phenomenon called serotype replacement, where vaccine strains are replaced by vaccine escape clones ^59^. A recent study showed that distinctive pneumococcal lineages exhibited same non-vaccine serotypes and dissimilar antibiotic resistance profiles^60^. Taken together, the evident replacement of serotypes (capsule genes transfer or expression) seems to have no association with the acquisition of macrolide resistance as observed in clinical strains and in our *ex vivo* bioreactor experiments.

## Material and Methods

### Bacterial strains, culture media, and antibiotics

In total 41 clinical isolated multidrug resistant (MIC of penicillin ≥ 8 µg/ml, erythromycin ≥128 µg/ml) *S. pneumoniae* strains were sequenced and used for *in silico* multilocus sequence typing (MLST), phylogenic tree construction, and recombination prediction. Serotyping of those clinical isolated were carried out using Latex and Quellung reaction (Statens Serum Institute, Copenhagen, Denmark). Pneumococcal strains were used for biofilm formation and antibiotic resistance transfer experiments are listed in Table 1. Strains were routinely cultured on blood agar plates (BAP), or grown in Todd Hewitt broth containing 0.5% (w/v) yeast extract (THY), at 37°C with a 5% CO2 atmosphere. Where indicated, streptomycin (200 µg/ml), trimethoprim (10 µg/ml), tetracycline (1 µg/ml), or/and erythromycin (1 µg/ml) was added to BAP. All antibiotics were purchased from Merck (Darmstadt, Germany).

### DNA Extraction and whole genome sequencing analysis

Genomic DNA of all tested pneumococcal isolates was extracted using a DNA mini kit (Qiagen, Valencia, CA, USA) and sent to whole genome sequencing (WGS) utilizing the Illumina HiSeq2000^TM^ platform. Sequence reads (Accession: PRJNA795524) were the mapped to a reference strain 19A-19339 (Accession: CP071917) to make single nucleotide polymorphism (SNP) calls using Snippy (v4.4.5) ^61^ and recombination prediction was then conducted via Gubbins (v2.4.1) ^62^ with the minimum number of 3 for base substitutions required to identify a recombination event. A paired-end fastq file for each tested strain was assembled by Shovill ^63^, with a minimal length and coverage of 200 bp and10x, respectively. The assemblies of selected recombinants were used to mapping against the complete sequence of strain D39 (NC_008533) in BRIG 0.95 ^64^.

D39 transformants were whole-genome sequenced using the NextSeq500 platform, targeting an average of 20M reads/sample (50X coverage) for the captured samples. The raw Illumina sequence reads were quality-tested by FastQC and trimmed by Trim Galore. Paired-end FastQ files were assembled using SPAdes (v3.11.1). The assembled capsule switch strain was annotated onto the closed genome sequence of the reference strain D39 (CP000410) and D39^Tn*17545*^ against GA17545 (AFGA01000000) using the online web-based RAPT and Genome Assembly Service tools. Annotated whole-genome sequences have been deposited in NCBI GenBank under BioProject no. PRJNA1055520.

### Inoculum preparation

The inoculum was prepared as previously described ^17^. Briefly, an overnight BAP culture was used to prepare a cell suspension in THY broth to an OD_600_ of ∼0.08. This suspension was incubated at 37°C in a 5% CO_2_ atmosphere until the culture reached an OD_600_ of ∼0.2 (early log phase), and then glycerol was added to a final 10% (vol/vol) and stored at −80°C until used. A frozen aliquot from each batch was removed to obtain the density (cfu/ml) by serial dilution and platting.

### Cell cultures

Human pharyngeal Detroit 562 cells (ATCC CCL-138) were cultured in Gibco™ Minimum Essential Media (MEM) (Thermo Fisher Scientific, Waltham, MA) supplemented with 10% non-heat-inactivated fetal bovine serum (FBS) (Atlanta biologicals), 1% nonessential amino acids (Sigma), 1% glutamine (Sigma), penicillin (10,000 U/ml)-streptomycin (10,000 µg/ml), and HEPES (10 mM) (Gibco). For the seeding, cells were harvested with 0.25% trypsin (Gibco), resuspended in the cell culture medium at a ratio of 1:5 and incubated at 37°C in a 5% CO_2_ humidified atmosphere.

### Bioreactor system

Detroit 562 cells were grown on Snapwell^TM^ filters (Corning, USA); these filters have a polyester membrane (0.4 µm) supported by a detachable ring. Once polarized (7-9 days), Snapwell-containing pharyngeal cells were immediately placed in a sterile vertical diffusion chamber (i.e., bioreactor) ^17^. Bioreactor chambers were perfused as detailed in our previous publication ^15^. Chambers were then inoculated with ∼1×10^8^ cfu/ml of each pneumococcal strain and incubated at ∼34°C under a sterile environment. At the end of the incubation Snapwell inserts were removed and pneumococci attached to human cells were washed once with PBS to remove planktonic cells. These pneumococci were harvested by sonication for 15 s in a Bransonic ultrasonic water bath (Branson, Danbury, CT), followed by extensive pipetting to remove all attached bacteria. An aliquot was used to obtain the density of each strain in the biofilm consortium, by serial dilution and platting on BAP containing the appropriate antibiotic, and another aliquot was plated onto BAP containing two, or three, antibiotics, to harvest recombinants.

### Confocal analysis of nasopharyngeal pneumococcal biofilm consortia

To visualize biofilm consortia by super-resolution and confocal microscopy, we installed a glass coverslip inside the Snapwell^TM^ filters prior to seeding human pharyngeal cells. Once pharyngeal cells became polarized, the Snapwell was installed in the bioreactor and inoculated as above. At the end of the incubation, the coverslip containing pharyngeal cells with pneumococcal biofilms were washed twice with PBS and fixed with 2% PFA for 15 min at room temperature. Once the fixative agent was removed, biofilms were washed with PBS and blocked with 2% bovine serum albumin (BSA) for 1 h at room temperature. These cells containing biofilms were simultaneously incubated with serotype-specific polyclonal antibodies (Statens Serum Institute, Denmark) (∼ 40 μg/ml), and wheat germ agglutinin (WGA) conjugated with Alexa Fluor (488 or 555, Invitrogen), for 1 h at room temperature. Antibodies had been previously labeled with Alexa-488 (anti-S4/A488) or Alexa-555 (anti-S2/A555 and anti-S6B/A555) following the manufacturer recommendations (Molecular Probes)^15, 18^. Stained preparations were finally washed two times with PBS and mounted with ProLong Diamond Antifade mountant with DAPI (Molecular Probes). Super-resolution confocal images were obtained using an Olympus FV1000 confocal microscope. Confocal micrographs were analyzed with ImageJ version 1.49k (National Institutes of Health, USA) or with the Imaris software 10.1.0 (Bitplane AG).

### Electron microscopy studies

Electron microscopy was used to visualize the detailed spatial localization of pneumococcal strains within nasopharyngeal biofilm consortia and their interactions. Instead of a glass coverslip as described above, we installed a square chip inside the Snapwell^TM^, or a Thermanox coverslip (Fisher scientific), and then cells were grown and then infected as detailed in the previous section. At the end of incubation, the square silicon chip, or Thermanox coverlis, was removed from the bioreactor culture chamber and processed as detailed below. Silicon chips, or Thermanox, were immediately fixed with a 2.5% glutaraldehyde solution in 0.1 M cacodylate buffer pH 7.4 overnight and then washed with the same buffer. Preparations were processed for scanning electron microscopy (SEM) as follows: post fixed with 1% osmium tetroxide and 1.5% potassium ferrocyanide in 0.1 M cacodylate buffer for one hour. They were subsequently rinsed with de-ionized water, followed by dehydration through an ethanol series ending with three exchanges of absolute ethanol. The samples were then placed into individual ventilated processing vessels in fresh absolute ethanol and placed into a Polaron E3000 critical point drying unit where the ethanol was exchanged for liquid CO_2_. This liquid CO_2_ was eventually brought to its critical point of 1073 psi at 31°C and allowed to slowly vent. Dried samples were then secured to labeled aluminum SEM stubs and coated with approximately 15 nm of chromium with a Denton DV-602 Turbo Magnetron Sputter coater. Samples were then viewed with a Topcon DS130F field emission scanning electron microscope using 5 kV accelerating voltage. Transmission electron microscopy (TEM). Preparations on Thermanox were infiltrated and embedded in Eponate 12 resin. Ultrathin sections were cut on a RMC PowerTome XL ultramicrotome at 70 nm, stained with 5% aqueous uranyl acetate and 2% lead citrate, and examined on a JEOL IEM-1400 transmission electron microscope equipped with Gatan UltraScan US1000.894 and Orius SC1000.832 CCD cameras.

### PCR studies of transformants

#### DNA Extraction and serotype-specific qPCR reactions

DNA was extracted from 200 μl of a fresh suspension of pneumococcal strains or a pool of recovered recombinants with the QIAamp DNA Minikit according to the manufacturer’s instructions. Final elution was done with 100 μl of elution buffer, DNA preps were quantified using a Nanodrop^TM^2000 spectrophotometer (Thermo Fisher, Wilmington, Delaware, USA). DNA preps were utilized as template for serotype-specific quantitative PCR reactions with primers and probes listed in Table 2 to identify the serotype of each strain as well as recombinants as detailed in our previous studies^15, 47, 65^.

#### Insertion of *ermB* near the capsule locus of strain TIGR4

The *ermB* gene was PCR amplified using DNA template from SPJV10^17^ and primers Ery-L-XbaI and Ery-R-XhoI listed in^17^. This *ermB* PCR product was purified using the QIAquick PCR Purification Kit (Qiagen, Valencia CA) and then digested with restriction enzyme XbaI. Simultaneously two ∼1.5 kb PCR products, to be cloned flanking *ermB,* were PCR amplified using DNA from strain TIGR4 as a template and primers JVS95L and JVS96R, to generate an upstream fragment, or JVS97L and JVS98R to generate a downstream. PCR products were purified as above and digested with XbaI (upstream) or XhoI (downstream), respectively. The XbaI-digested upstream fragment was first ligated using T4 DNA ligase and the ligated product used as a template in PCR reactions using primers JVS95L and Ery-R-XhoI. This purified PCR product was digested with XhoI and ligated to the XhoI-digested downstream fragment as mentioned. The final ligated fragment was used as a template in PCR reactions with primers JVS95L and JVS98R, purified as indicated above and sequenced at Eurofins to confirm the construct. This cassette containing sequences downstream the capsule locus was transformed (∼100 ng) into competent cells of strain TIGR4 wt. A transformant recovered in BAP with erythromycin (1 µg/ml), named SPJV24, was serotyped with Quellung antisera (Statens Serum Institute, Copenhagen, Denmark) to confirm expression of serotype 4 capsule, and then DNA extracted to confirm the *ermB* was located upstream the capsule locus, as detailed in the Results section.

#### Construction of a TIGR4^ΩermB^Δ*comCDE* mutant

The mutation was prepared by replacing the operon *comCDE* in TIGR4^ΩermB^ with a truncated fragment containing the *catP* gene. This fragment was amplified from a GA16833Δ*comCDE* mutant (SPJV37) by PCR with primers JVS0413 and JVS0414 (Table 3). The mutants were selected on BAP containing chloramphenicol (3 µg/ml). The mutation was confirmed by PCR in the resulting chloramphenicol-resistant clones.

**Table 3.**
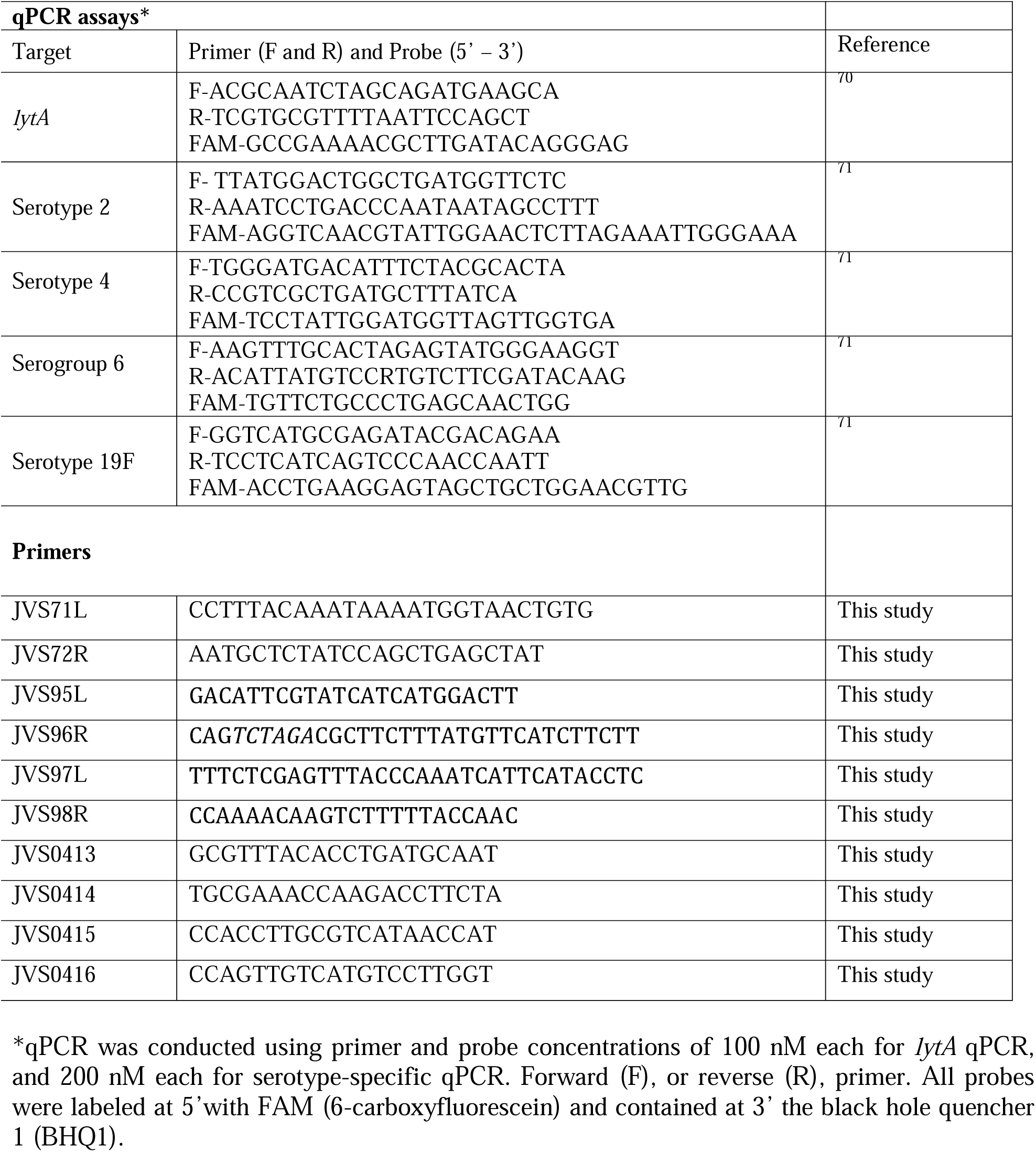
Quantitative PCR assays and primers utilized in this study.

#### Primary metabolism analysis of mixtures of *S. pneumoniae* strains infecting human pharyngeal cells

D39 and GA16833 or D39Δ*comD* and GA16833, were infected into the bioreactor as previous detailed, and infected cells were incubated at 37°C. The influx coming off the bioreactor chamber (i.e., supernatant and planktonic bacteria) was collected for 60 min after the first h of incubation, or after 3 h of incubation. Supernatants were filter sterilized with a 0.4 μm syringe filter and immediately frozen at −80°C. Primary metabolism by gas chromatography-time of flight mass spectrometry (GC-TOF MS) was performed at the West Coast Metabolomics Center, UC Davis. Briefly, samples extracted using 1ml of 3:3:2 ACN:IPA:H_2_O (v/v/v). Half of the sample was dried to completeness and then derivatized using 10 μl of 40 mg/ml of methoxyamine in pyridine. They were shaken at 30°C for 1.5 h. Then 91 μl of MSTFA + FAMEs to each sample and they were shaken at 37°C for 0.5 h to finish derivatization. Samples were then vialed, capped, and injected onto the instrument. We use a 7890A GC coupled with a LECO TOF. 0.5 μl of derivatized sample is injected using a splitless method onto a RESTEK RTX-5SIL MS column with an Intergra-Guard at 275°C with a helium flow of 1 ml/min. The GC oven is set to hold at 50°C for 1 min then ramp to 20°C/min to 330°C and then hold for 5 min. The transferline was set to 280°C while the EI ion source was set to 250°C. The Mass spec parameters collect data from 85m/z to 500m/z at an acquisition rate of 17 spectra/sec. Raw data were processed by LECO ChromaTOF version 4.5 for baseline subtraction, deconvolution and peak detection, while BinBase was used for annotation and reporting^66^.

### Statistical analysis

We performed one-way analysis of variances (ANOVA) followed by Dunnet’s multiple comparison test, when more than two groups were compared or the Student *t* test to compare two groups, as indicated. All statistical analysis was performed using the software Graph Pad Prism (version 8.3.1).

## Acknowledgements

This study was primarily supported by National Institutes of Health (NIH; R21AI112768-01A1), the National Natural Science Foundation of China (No. 32000092) to XW, and the Jinhua Science and Technology Research Key program (No. 2021-3-07) to XX. JEV is also supported by a grant from NIGMS through the Molecular Center of Health and Disease (1P20GM144041-01A1 7651). The work performed through the UMMC Molecular and Genomics Facility is supported, in part, by funds from the NIGMS, including the Mississippi INBRE (P20GM103476) and Obesity, Cardiorenal and Metabolic Diseases-COBRE (P30GM149404). Special thanks to Dr. Lesley McGee and Dr. Bernard Beall from the Centers for Disease Control and Prevention (CDC) for providing pneumococcal invasive isolates, and to Dr Shanshan Zhao for provide clinical multidrug resistant pneumococci. We thank Dr. Hong Yi, and Dr. Jeannette Taylor for their assistance with electron microscopy at the Robert P. Apkarian Integrated Electron Microscopy Core in Emory University. Authors appreciate the assistance of Dr. Neil Anthony, from Emory University School of Medicine, with confocal microscopy, and Dr. Veronique Parrot for her assistance on preparing some figures.

